# Broad-scale mercury bioaccumulation patterns in two freshwater sport fishes: testing the role of growth dilution in a warming climate

**DOI:** 10.1101/280107

**Authors:** Shyam M. Thomas, Stephanie J. Melles, Satyendra P. Bhavsar

**Affiliations:** Department of Chemistry & Biology, Ryerson University, Toronto; Ontario Ministry of the Environment and Climate Change, Toronto

**Keywords:** fish growth rates, mixed-effects models, Northern Pike, Ontario, spatiotemporal trends, temperature effects, Walleye

## Abstract

Sport fishes at the apex of aquatic food webs are indicators of mercury in the environment. However bioaccumulation of mercury in fish is a complex process that varies in space and time. Both large-scale climatic and environmental, as well as biological factors are drivers of these space-time variations. In this study, we avail a long-running monitoring program from Ontario, Canada to better understand spatiotemporal variations in fish mercury bioaccumulation. Focussing on two common large-bodied fishes (Walleye and Northern Pike), the data were first stratified by latitudinal zone (north, mid, and south) and eight temporal periods (between 1975 & 2015). A series of linear mixed-effects models (LMEMs) with latitudinal zone, time period, and their interactions as random effects were used to capture the spatial, temporal, and spatiotemporal variations in mercury bioaccumulation. The random slopes from the best-fitting LMEM were used to define bioaccumulation index and capture trends in space and time. Given the generally warming climate trend over the past 45 years, the role of growth dilution in modulating the bioaccumulation trends was also evaluated. The full model comprising of space, time and space-time interactions was the best-fit with interaction effects explaining most of the variation. Spatiotemporal trends showed overall similar patterns for both species. Growth dilution in conjunction with estimated rates of warming for different latitudinal zones failed to explain the spatiotemporal trends. Temporal trends showed contrasting bioaccumulation patterns-increasing in Northern Pike and decreasing in Walleye, suggesting temperature-driven growth dilution is more likely in latter. However, a space-for-time substitution revealed only a weak presence of growth dilution in Walleye, and it was not attributable to temperature differences. Overall, our study summarizes broad-scale variations in fish mercury and explores the role of growth dilution in shaping the observed patterns.

## Introduction

Mercury pollution captured global attention during the tragic Minamata poisoning, which highlighted the fatal neurotoxic effects of consuming mercury tainted fish. Despite concerted global efforts to reduce mercury emissions, mercury as an anthropogenic pollutant remains a matter of grave concern, since in elemental form mercury is highly mobile, and often ends up in aquatic and terrestrial ecosystems far from the emission source (Driscoll et al 2013). In aquatic ecosystems the impact of mercury pollution is mostly associated with organic methylmercury (MeHg). MeHg is all the more detrimental as it can biomagnify and bioaccumulate such that older large-sized fishes at the apex of aquatic food chain have disproportionately higher amounts of mercury (Driscoll et al 2013). In short, large-bodied (sport) fishes at the top of food chains are good indicators of mercury levels, and when consumed by humans can adversely affect human health.

Both the United States and Canada have established monitoring programs at state and provincial levels to study fish mercury dynamics. Analyses of these large-scale databases typically show a declining trend in fish mercury levels between 1970 and 2000 (Chalmers et al. 2011; Monson et al. 2011; Gandhi et al. 2014). However, there is substantial geographic variation in these long-term trends (Kamman et al. 2005; Chalmers et al. 2011; Gandhi et al. 2014). Most importantly, many regions are experiencing increasing mercury levels in several key sport fishes in recent years, and this may have severe implications for human health if the trend continues (e.g., Gandhi et al. 2014, Gandhi et al. 2015). Climate change, particularly the rapidly warming climatic conditions experienced by lakes in temperate regions is thought to be one of the likely reasons for the recent surge in mercury levels (e.g., Gandhi et al. 2014, Chen et al. 2018). Warming climate can affect fish mercury levels via several processes operating at different scales, which are poorly understood to date. At the lake and watershed level, warming conditions are known to increase net methylation rates, and thus increase the overall amount of bioavailable methylmercury in a lake (Canário et al. 2007; Stern et al. 2011). Warmer temperatures affects mercury levels at the community level too, by altering trophic position, food-chain length, and productivity (Kidd et al. 2012; Lavoie et al. 2013). However, these impacts are highly variable and are contingent on the species of fish and species composition (Stern et al. 2011). Ecosystem and community level effects together determine the amount of bioavailable MeHg, which sets the exposure baseline.

The amount of mercury in a fish is eventually determined by consumption of mercury contaminated food at the individual level, and the final amount retained is a complex function of growth, consumption, and metabolism. Fish with better growth efficiency (i.e., ratio of consumption rate to growth rate) tend to accumulate less mercury, while increased metabolism may demand higher rates of consumption thus increasing accumulation of mercury (Ward et al. 2011; Dijkstra et al 2013). Fast growing fish with greater growth efficiency are hypothesized to accumulate less mercury since the net amount of biomass added is much greater for every unit of mercury gained, and this is referred to as growth dilution (GD). Studies also suggest that GD is most likely in situations where fast growth rate is accompanied with low metabolic costs or high quality food (low in mercury contamination) (Karimi et al. 2007; Ward et al. 2010; Dijkstra et al 2013). Rising temperatures are expected to boost fish growth rates and resultant GD as long as metabolic costs remain below species threshold levels, and these processes could potentially reduce overall fish mercury levels. However, warmer temperatures may also increase metabolic costs thus potentially offsetting any reduction in fish mercury levels achieved via increased growth rates. As mentioned earlier, warmer temperatures can increase the amount of MeHg through enhanced microbial activity or increase in food-chain length, which eventually can affect fish mercury levels. Put together, rising temperature can impact fish mercury levels in a complex manner. Not surprisingly, few studies have explored the role of warming climate on fish mercury levels (Dijkstra et al 2013). Moreover, studies that have found evidence of GD as a key modulating mechanism of mercury levels are either experimental studies or based on observations from few lakes without any consideration of large-scale climatic factors. This particular paucity of studies on growth dilution in the context of warming climate is not surprising, since large-scale databases with fish age information are scarce.

Mercury bioaccumulation also implies mercury concentration in fish varies strongly with fish size and age (Gewurtz et al. 2011a,b). Understandably, ecological studies on fish mercury are usually based on a standard fish size or age class defined around the sample median (Kamman et al. 2005; Gewurtz et al. 2011a). Similarly, studies describing historical and contemporary trends in fish mercury levels from monitoring data are also usually based on variations within similar standardized size classes (Chalmers et al. 2011; Gandhi et al. 2014). This practice of using standardized size or age classes in fish mercury studies is performed to account for the influence of exposure time and thereby to minimize variability in mercury concentrations. However, this approach can potentially lead to substantial information loss as variations outside the standard size classes are ignored, which can result in ambiguous estimates of fish mercury trends.

In this study, we first develop a bioaccumulation index based on full range of variation in fish size and mercury levels for two common freshwater sport fishes. To do so, we make use measurements from Ontario’s long running monitoring program with data spanning 15 latitudinal degrees and 45 years. The bioaccumulation index is then used to characterize the large-scale mercury bioaccumulation trends in space and time. We further explore the summarized spatiotemporal trends in mercury bioaccumulation by explicitly considering GD effects within a climate change context. In the second part of our study, we take advantage of another Ontario-wide monitoring dataset of comparable spatial scope that includes fish age information. Using the age information and a space-for-time latitudinal substitution approach, we test the role of growth dilution in explaining the observed species-specific temporal trends in mercury bioaccumulation.

## Methods

### Fish Mercury Data

We used one of the largest known fish monitoring databases compiled by the Ontario Ministry of the Environment and Climate Change (OMOECC) that tracks pollutant loads in several key sport fishes. The Ontario-wide monitoring program began in 1970, i.e. a temporal scope of more than 45 years (1970 onwards) and covers a broad climatic range with a latitudinal breadth of nearly 15 degrees (41.5° to 56.5°). Each data record provides fish species identity, length, body mass, sex and amount of mercury alongside information on lake or waterbody identity/name, and geo-coordinates of the location where the fish was sampled. With multiple fish samples often taken at a given time and location and a total of 126,652 records, the database provides a comprehensive picture of fish mercury levels and several key fish-level attributes.

### Data Selection & Focal Species

In order to develop a bioaccumulation index that captures variation in mercury levels across a broad range of climatic conditions, we chose species with the most widespread distribution across Ontario. Walleye (*Sander vitreus*) and Northern Pike (*Esoxlucius*) were the best candidate species in this respect; with their broad nearly identical distribution patterns that span most of Ontario (Figure 1a &b). Walleye is a native cool-water predatory fish that is common in most lakes of Canada and northern United States. Walleye, like many shoaling fishes, prefers large open waters, and its large eyes enable it to hunt effectively in low light conditions, particularly at dusk and at night (Swenson 1977). Northern Pike is an equally common cool-water predatory fish with a broad pan-artic distribution that includes Europe, Russia, Canada and northern United States. Unlike Walleye, Northern Pike is a large ambush predator that prefers to hunt during the day and like most ambush predators they need cover in the form of dense vegetation or submerged logs (Casselman and Lewis 1996). Also notable is the difference in body size with Walleye typically being smaller in size than Northern Pike. In Ontario, Northern Pike are known to attain an average size of 45-75 cm, whereas Walleye typically range between 35-58 cm. Walleye and Northern Pike are consumers at the top of aquatic food-chains that co-occur in several lakes and freshwater bodies in Ontario, however their ecology, feeding habits and growth differ substantially, making this pair of sport fishes particularly interesting to detect species-specific differences in mercury bioaccumulation. In a final data selection process, fish samples collected during the first 5 years (i.e. 1970-1974) were not included as they were selectively collected from locations within close proximity to known sources of mercury pollution, and hence had disproportionately higher mercury levels. In summary, records between 1975 and 2015 of Walleye and Northern Pike were analyzed to develop the bioaccumulation index.

**Figure 1.**
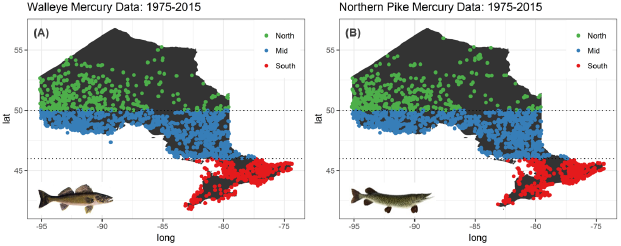
Sampling distribution of fish mercury samples in Ontario (1975-2015) for Walleye and Northern Pike across the defined latitudinal zones.

### Mercury Data Analyses

To capture large-scale spatiotemporal patterns, data of each species were divided into 8 temporal periods (1975-79,1980-84, 1985-90, 1990-94, 1995-99, 2000-04, 2005-09, 2010-15) and 3 latitudinal zones (south: 40°N – 46°N, mid: 46°N – 50°N, and north: >50°N). The temporal categories are essentially 5-year periods, except for the ‘2010-15’ period, while each latitudinal zone has a range of 5 latitudinal degrees. We analyzed fish mercury-body size relationships based on all possible combinations spatial and temporal categories, thus effectively capturing spatial, temporal, and spatiotemporal effects.

Mercury levels in fish typically follow an allometric relationship with body size (Gewurtz et al. 2011):

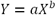

where *Y* is amount of mercury (*μg*/*g*) and *X* is fish body length (cm). When log-transformed, the allometric relationship takes the form of a linear model:

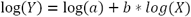

We use this log-transformed model for all analyses. Specifically, to quantify the effect of space, time, and space-time interactions, we make use of linear mixed-effects models (LMEM) with latitudinal zones and temporal periods as crossed random effects (Bolker et al. 2009). All LMEMs were fit using lme4 package (Bates et al. 2014) and further analyzed with merTools package (Knowles and Fredrick 2016) in R. In total, four distinct LMEM’s were separately fit to Walleye and Northern Pike data in order to analyze species-specific variations in mercury bioaccumulation. It may be noted that fish growth models are typically non-linear and they are often analysed using more complex non-linear models such as Gompertz and von Bertalanffy growth functions (Gamito 1998; Katsanevakis and Maravelias 2008). However, for easy interpretation of model parameters such that they effectively capture latitudinal variation in growth and mercury bioaccumulation, we assumed a linear relationship by log-transforming both response and predictor variables. All model comparisons were done using Akaike Information Criterion (AIC), where lower AIC values imply better model fit. The four models can be summarized as shown below:

a. Spatial Effects:

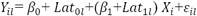
b. Temporal Effects:

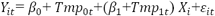
c. Spatial and Temporal Effects:

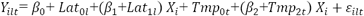
d. Spatiotemporal Effects:

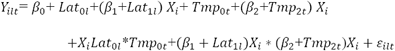

where, *Y*_*ilt*_ represents mercury concentration found in fish *i* collected at latitudinal zone *l* during time period *t*, and *X* implies fish length, which is the predictor variable. *Lat* and *Tmp* are latitudinal zones and temporal periods, respectively, which are the random effects in the LMEMs. The parameters inside the parenthesis together form the random slope coefficients, which indicate the magnitude of bioaccumulation for a given latitudinal zone *l*, and temporal period *t*. These random slope coefficients when compiled together form a *‘bioaccumulation index’* for each unique set of random factors and their combination (i.e. spatial, temporal, and spatiotemporal effects). In summary, for each fish species the bioaccumulation index derived from the LMEMs describes variation in mercury bioaccumulation across either latitudinal zones, temporal periods, or a combination of latitudinal zones and temporal periods.

### Climate Data

To provide an environmental context for the long-term bioaccumulation trends, climatic conditions in Ontario were summarized for 45 years (1970-2015) using temperature, growing degree-days and precipitation measure. Specifically stated, the reason for including broad-scale climatic trends was to deduce the potential role of growth dilution in modulating fish mercury bioaccumulation. The data were sourced from Environment Canada’s historical climate data website (http://climate.weather.gc.ca/index_e.html). The website provides climate information for each station at daily, monthly, and annual intervals. As with spatiotemporal trends of bioaccumulation, climate trends were captured across 5-year periods for each of the three latitudinal zones. There were a total of 9 temporal periods with the earliest being 1970-75 and ending at 2010-15. Within each latitudinal zone, stations with complete climate data were used to characterize the climate trends. However for many stations complete climate data spanning the entire 45-year period were not available, which resulted in the selection of very few compatible stations. Thus, three weather stations each were selected for south (Trenton, Ottawa and Glasgow) and mid (Chalk, Sudbury and Kenora) latitudinal zones, while for north only two weather stations were available (Sioux and Moosonee). In summary, average daily temperature and precipitation were estimated for each 5 year period and latitudinal zones, while number of growing degree days were estimated as the cumulative number of days when average daily temperature was above 5 °C.

### Role of growth dilution

The long-term (40 years) nature of fish mercury data implies both spatiotemporal and temporal trends in mercury bioaccumulation are likely to be influenced by changing climatic conditions, particularly given the increasing temperature conditions. To deduce this, we first explored LMEM-derived bioaccumulation indices to see if the rate of change in temperature and growing-degree days for the three different latitudinal zones explain observed spatiotemporal bioaccumulation trends in Walleye and Northern Pike. We hypothesized that warmer temperatures are likely to increase fish growth rates resulting in increased growth dilution and eventually leading to overall decreases in bio-accumulated mercury levels. Thus, if growth dilution due to temperature driven variation in fish growth rates is the primary mechanism, then latitudinal zones experiencing the greatest rate of increase in temperature conditions are likely to show the sharpest decline in mercury levels compared to latitudinal zones experiencing relatively modest rates of increase. Unlike spatiotemporal trends in mercury bioaccumulation, temporal trends without any spatial variation capture mercury bioaccumulation pattern during the 40-year warming period. We further examined whether the observed temporal bioaccumulation trends in Walleye and Northern Pike were explained by growth dilution in a warming climate. Specifically, we hypothesized that if growth dilution is due to temperature dependent variation in growth rates, then species that show increase in growth rate with temperature will show a decreasing trend in mercury bioaccumulation with increasing temperature conditions as warmer temperatures are expected to strengthen growth dilution. On the other hand, species that show decrease in growth rate with temperature will yield an increasing bioaccumulation trend with increasing temperature as warmer conditions are now expected to weaken the effect of growth dilution. In order to detect this species-specific difference in growth dilution, estimation of growth rate based on fish age is necessary. To this end, we made use of another database with age information – the broad-scale monitoring (BsM) database. The BsM program was developed largely to standardize data collection and manage fisheries at broad-scales for the entire Province of Ontario by sampling a representative number of lakes every 5 years. The first such sampling cycle covered the years 2008-2012, which we avail to estimate growth rates of Walleye and Northern Pike.

Next we used a space-for-time substitution approach, wherein BsM data covering Ontario were first used to capture latitudinal variation in both growth rates and mean mercury levels. The latitudinal variation captured the underlying difference in growth dilution due to varying temperature conditions, since latitude and temperature show strong inverse correlation. Specifically stated, the broad latitudinal coverage of BsM data spans temperature conditions that range from warm southern lakes to cold northern lakes, and this apparent temperature gradient serves as a template to test the role of variation in growth dilution, which can then be substituted for time to explain temporal trends (see Figure 2 for details). We chose a space-for-time substitution approach over a more direct estimate of temporal variation in growth rate because latitudinal gradient captures temperature differences more substantively and consistently than a temporal sequence of years, thus providing a stronger basis to test the role of temperature-driven variation in fish growth rates. Latitudinal variation in growth rates were estimated using linear mixed-effects models (LMEMs) with latitudes as random effects, such that the random slopes are latitude-specific estimates of growth rate. In summary, the additional LMEM’s were fitted to Walleye and Northern Pike data to capture their growth rate.

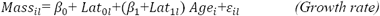

where, *Mass*_*il*_ represents body mass of fish, *i* collected at latitude, *l*, *Age* (fish age) is the predictor variable in the growth model, and *Lat* represents all unique latitudes as random effects. Finally, presence of growth dilution was tested by running a correlation analysis between the latitude-specific estimates of mean mercury levels and growth rates. A negative slope indicates presence of growth dilution, while the correlation coefficient (Pearson’s r) states how well growth rates explained the variation in mercury levels.

It may be noted that unlike in the analyses of mercury bioaccumulation where fish length was used to capture trends in space and time, fish mass is used here in the analysis of growth dilution. We maintain this distinction for three reasons: 1) mercury levels in fish are typically reported using fish length as the primary covariate and most studies on mercury bioaccumulation trends are based on standard fish length, 2) growth rates are sensitive to fish length as a predictor, especially when comparing growth rates and consequent growth dilution between fish species (i.e., Walleye & Northern Pike) with distinct body forms (Tom Johnston *personal communication*), and 3) fish length and mass are highly correlated with R^2^ > 0.9 for both species (Supplementary Figure S1), thus effectively allowing either to be used as a proxy for the other

## Results

**T**he long-term monitoring program spanning 40 years (1975-2015) resulted in 49,690 Walleye samples and 32,636 Northern Pike samples. Among the four models of mercury bioaccumulation, the full model that combined spatial, temporal, and spatiotemporal interactions was the best fit with lowest AIC values for both Walleye and Northern Pike (Table 1a &b). Models of spatial effects and temporal effects alone had poor fits and high AIC values, whereas the model with spatial and temporal effects together had a better fit with lower AIC values. Within the best-fitting full model, much of the large-scale variation in mercury levels was captured by the random grouping factor representing space-time interaction effects. This is evident from the substantially larger intra-cluster correlation coefficient (ICC) estimates of 0.735 and 0.672 for Walleye and Northern Pike, respectively (*i.e.* ICC_LatZones:Period_ in Table 1a &b). In mixed effects models where data are typically divided into different clusters or groups, ICC describes the amount of variance explained by a grouping factor relative to the total variance explained by all grouping factors involved and amount of residual within-group variance.

**Table 1a.** Summary statistics of the four different LMEMs used to characterize spatial, temporal, and spatiotemporal effects in Walleye the data stratified into three latitudinal zones (LatZones) and eight temporal periods (Period).

|  | *Spatial* |  |  | *Temporal* |  |  | *Space & Time* |  |  | *Full* |  |  |
| --- | --- | --- | --- | --- | --- | --- | --- | --- | --- | --- | --- | --- |
|  | *B* | *CI* | *p* | *B* | *CI* | *p* | *B* | *CI* | *P* | *B* | *CI* | *p* |
| *Fixed Parts* |  |  |  |  |  |  |  |  |  |  |  |  |
| Intercept | -6.54 | -6.88 – -6.20 | <.001 | -6.23 | -6.62 – -5.84 | <.001 | -6.62 | -6.99 – -6.25 | <.001 | -6.8 | -7.47 – -6.13 | <.001 |
| log(length) | 1.54 | 1.50 – 1.59 | <.001 | 1.44 | 1.32 – 1.55 | <.001 | 1.55 | 1.49 – 1.62 | <.001 | 1.6 | 1.45 – 1.74 | <.001 |
| *Random Parts* |  |  |  |  |  |  |  |  |  |  |  |  |
| $\sigma^2$ | | 0.488 | | | 0.522 | | | 0.458 | | | 0.442 | |
| $\tau_{00, \text{LatZones:Period}}$ | | --- | | | --- | | | --- | | | 1.648 | |
| $\tau_{00, \text{Period}}$ | | --- | | | 0.301 | | | 0.063 | | | 0.019 | |
| $\tau_{00, \text{LatZones}}$ | | 0.085 | | | --- | | | 0.077 | | | 0.133 | |
| $\rho_{01}$ | | 0.604 | | | -0.943 | | | -0.849 | | | -0.993 | |
| $N_{\text{LatZones:Period}}$ | | --- | | | --- | | | --- | | | 24 | |
| $N_{\text{Period}}$ | | --- | | | 8 | | | 8 | | | 8 | |
| $N_{\text{LatZones}}$ | | 3 | | | --- | | | 3 | | | 3 | |
| $\text{ICC}_{\text{LatZones:Period}}$ | | --- | | | --- | | | --- | | | 0.735 | |
| $\text{ICC}_{\text{Period}}$ | | --- | | | 0.366 | | | 0.105 | | | 0.008 | |
| $\text{ICC}_{\text{LatZones}}$ | | 0.149 | | | | | | 0.129 | | | 0.059 | |
| Observations |  | 49690 |  |  | 49690 |  |  | 49690 |  |  | 49690 |  |
| $R^2$ | | .316 | | | .269 | | | .358 | | | .381 | |
| AIC |  | 105392.353 |  |  | 108774.404 |  |  | 102283.674 |  |  | 100658.545 |  |

Abbreviations and Symbols: $\sigma^2$ = residual (within-group) variance; $\tau_{00}$ = between-group variance; $\rho_{01}$ = correlation between random slope and random intercept; $N$ = number of groups; ICC = intra-class correlation coefficient (see Methods for more details); $R^2$ = r-squared value; AIC = Akaike Information Criterion

**Table 1b.** Summary statistics of the four different LMEMs used to characterize spatial, temporal, and spatiotemporal effects in Northern Pike with the data grouped into three latitudinal zones (LatZones) and eight temporal periods (Period).

|  | *Spatial* |  |  | *Temporal* |  |  | *Space & Time* |  |  | *Full* |  |  |
| --- | --- | --- | --- | --- | --- | --- | --- | --- | --- | --- | --- | --- |
|  | *B* | *CI* | *p* | *B* | *CI* | *p* | *B* | *CI* | *p* | *B* | *CI* | *p* |
| *Fixed Parts* |  |  |  |  |  |  |  |  |  |  |  |  |
| Intercept | -6.45 | -7.9 – -5 | <.001 | -7.33 | -8.27 – -6.40 | <.001 | -7.21 | -8.4 – -5.9 | <.001 | -7.63 | -8.3 – -6.8 | <.001 |
| log(length) | 1.41 | 1.1 – 1.7 | <.001 | 1.61 | 1.40 – 1.82 | <.001 | 1.58 | 1.34 – 1.82 | <.001 | 1.68 | 1.5 – 1.8 | <.001 |
| *Random Parts* |  |  |  |  |  |  |  |  |  |  |  |  |
| $\sigma^2$ | | 0.500 | | | 0.515 | | | 0.474 | | | 0.458 | |
| $\tau_{00, \text{LatZones:Period}}$ | | --- | | | --- | | | --- | | | 1.897 | |
| $\tau_{00, \text{Period}}$ | | --- | | | 1.789 | | | 1.024 | | | 0.471 | |
| $\tau_{00, \text{LatZones}}$ | | 1.704 | | | --- | | | 0.767 | | | 0.000 | |
| $\rho_{01}$ | | -0.998 | | | -0.996 | | | -0.993 | | | -0.996 | |
| $N_{\text{LatZones:Period}}$ | | --- | | | --- | | | --- | | | 24 | |
| $N_{\text{Period}}$ | | --- | | | 8 | | | 8 | | | 8 | |
| $N_{\text{LatZones}}$ | | 3 | | | --- | | | 3 | | | 3 | |
| $\text{ICC}_{\text{LatZones:Period}}$ | | --- | | | --- | | | --- | | | 0.672 | |
| $\text{ICC}_{\text{Period}}$ | | --- | | | 0.776 | | | 0.452 | | | 0.167 | |
| $\text{ICC}_{\text{LatZones}}$ | | 0.773 | | | --- | | | 0.339 | | | 0.000 | |
| Observations |  | 32636 |  |  | 32636 |  |  | 32636 |  |  | 32636 |  |
| $R^2$ | | .293 | | | .272 | | | .330 | | | .354 | |
| AIC |  | 70055.664 |  |  | 71066.828 |  |  | 68379.025 |  |  | 67314.081 |  |

Abbreviations and Symbols: Same as Table 1a

### Analysis of fish mercury in the context of climate

Estimates of random slopes from the full model, indicating the magnitude of bioaccumulation (i.e. bioaccumulation index in Tables 3a-c), showed distinct spatiotemporal patterns that were consistent for both Walleye and Northern Pike (Figures 3a & b). In Walleye, the south latitudinal zone showed strong decline in bioaccumulation with time (β = -0.026; R^2^ = 0.81), whereas the mid zone showed a relatively weak positive trend (β = 0.0083; R^2^ = 0.35) and the north latitude showed a very weak positive trend (β = 0.0045; R^2^ = 0.05). For Northern Pike, bioaccumulation increased strongly with time in north latitudes (β = 0.033; R^2^ = 0.74), while mid latitudes showed a subtle increase (β = 0.0056; R^2^ = 0.18) and south latitudes showed a general decline with time (β = -0.008; R^2^ = 0.19). Overall, both species seem to be bioaccumulating mercury at an increasing rate in the relatively colder latitudinal zones of mid and north, while in the warmer southern latitudes rate of bioaccumulation seems to be decreasing. Unlike spatiotemporal trends that appear similar overall for both Walleye and Northern Pike, a purely temporal perspective (i.e. time as the only random factor) highlights an interesting dissimilarity with contrasting trends in bioaccumulation: Walleye bioaccumulation trends declined over time, whereas Northern Pike showed a strong increase (Figure 3c). Unlike the bioaccumulation trends, climate variables showed an overall increasing trend (Figure 3d – f), and this was particularly consistent in the case of average daily temperature (β = 0.044; R^2^ = 0.96) and growing degree-days (β =6.3; R^2^ = 0.83). As expected, the overall average (i.e. the intercept) of annual mean temperatures and growing degree-days decreased with increase in latitude. However, it is worth noting that the rate of increase (i.e. the slope) was greatest in northern latitudes (β_temp_ = 0.06; β_gdd_ = 7.3) followed by mid (β_temp_ = 0.04; β_gdd_ = 5.9) and southern latitudes (β_temp_ = 0.034; β_gdd_ = 5.6), thus suggesting that northern latitudes are warming at a faster rate than southern latitudes. Average daily precipitation, like temperature and growing degree-days, showed a generally increasing trend with time, with northern latitudes recording highest rates of increase. However, there was substantial variation during the 45-year time period as evident from the generally lower R^2^ values of precipitation compared to those of temperature and growing degree-days. Also, average daily precipitation was on the whole greater in southern Ontario relative to mid and northern regions.

**Figure 2.**
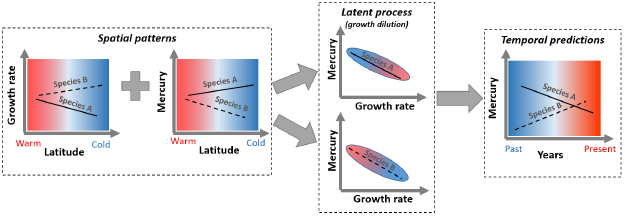
The space-for-time latitudinal substitution. Latitudinal variation in growth rate and mercury concentrations (left panel) can result in contrasting mercury bioaccumulation temporal trends (right panel) via two distinct temperature-driven growth dilution effects (middle panel) a highlighted by the hypothetical species A and B. In species A growth rate declines and mercury levels increases in colder conditions resulting in positive temperature driven growth dilution (i.e. growth dilution increases with increase in temperature), which when substituted in time results in a declining temporal trend wherein past years with colder temperatures have higher mercury levels relative to more recent times with warmer temperature conditions. On the other hand, species B with contrasting spatial patterns yields a negative temperature driven growth dilution (i.e. growth dilution increases with decrease in temperature), which when substituted in time results in an increasing temporal trend.

**Figure 3.**
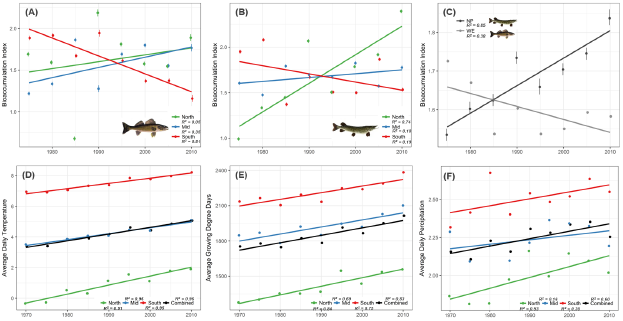
Spatiotemporal trends of mercury bioaccumulation in (a) Walleye and (b) Northern Pike as predicted by the random slopes of the full model with time period and latitudinal zone interactions as random effects. Temporal trends in bioaccumulation for both Walleye and Northern Pike are shown in c) as the predicted random slopes of the full model with time period alone as the random effect. Change in climatic conditions are shown as spatiotemporal trends in (d) temperature, (e) growing degree days, and (f) precipitation based on average measures at Environment Canada sampling stations, estimated from aggregated 5-year time periods.

### Latitudinal variation in growth rate

The BSM data overall comprised of 3159 Walleye and 1699 Northern Pike samples collected from lakes across Ontario ranging from 44.5° N to 55° N (Figures 4a,b). LMEMs testing for latitudinal variation in growth rate with fish age as the predictor yielded significant results for both Walleye and Northern Pike (Table 2). This is evident from the high ICC values associated with growth rate LMEMs. Moreover, latitudinal variation in growth rates highlighted interesting difference between the two species. Stated specifically, the random slopes describing latitude-specific growth rates showed a negative relationship with latitude in Walleye suggesting growth rate decreased with latitude, whereas the mean latitudinal mercury levels showed a weak positive correlation with latitude (Figures 5a,c). Northern Pike, on the other hand, showed a positive relationship between latitude and growth rates as well as between latitude and mean mercury suggesting both growth rates and mercury levels increased with latitude in Northern Pike (Figures 5b,d).

**Table 2.** Summary statistics of LMEMs describing latitudinal variation in growth for Walleye and Northern Pike with data grouped by unique latitudes (LAT).

|  | *Walleye* |  |  | *Northern Pike* |  |  |
| --- | --- | --- | --- | --- | --- | --- |
|  | *B* | *CI* | *p* | *B* | *CI* | *p* |
| **Fixed Parts** |  |  |  |  |  |  |
| Intercept | 4.05 | 3.92 – 4.18 | <.001 | 5.44 | 5.36 – 5.53 | <.001 |
| Log(Fish Age) | 1.34 | 1.27 – 1.40 | <.001 | 1.05 | 1.00 – 1.09 | <.001 |
| **Random Parts** |  |  |  |  |  |  |
| $\sigma^2$ | | 0.092 | | | 0.087 | |
| $\tau_{00, LAT}$ | | 0.882 | | | 0.215 | |
| $\rho_{01}$ | | -0.896 | | | -0.771 | |
| $N_{LAT}$ | | 291 | | | 240 | |
| $ICC_{LAT}$ | | 0.906 | | | 0.713 | |
| Observations |  | 3159 |  |  | 1699 |  |
| $R^2$ | | .874 | | | .847 | |

Abbreviations and Symbols: $\sigma^2$ = residual (within-group) variance; $\tau_{00}$ = between-group variance; $\rho_{01}$ = correlation between random slope and random intercept; $N_{LAT}$ = number of groups (unique latitudes); $ICC$ = intra-class correlation coefficient (see Methods for more details); $R^2$ = r-squared value; $AIC$ = Akaike Information Criterion.

**Figure 4.**
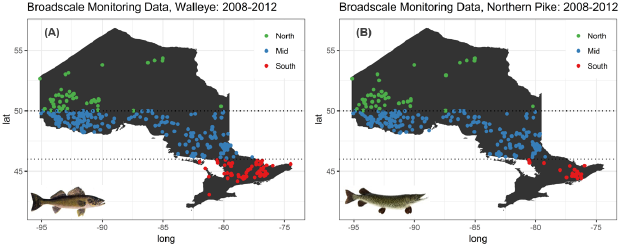
Distribution of Walleye and Northern Pike samples in Ontario across the three defined latitudinal zones, obtained from the Broad Scale Monitoring program’s first sampling cycle.

**Figure 5.**
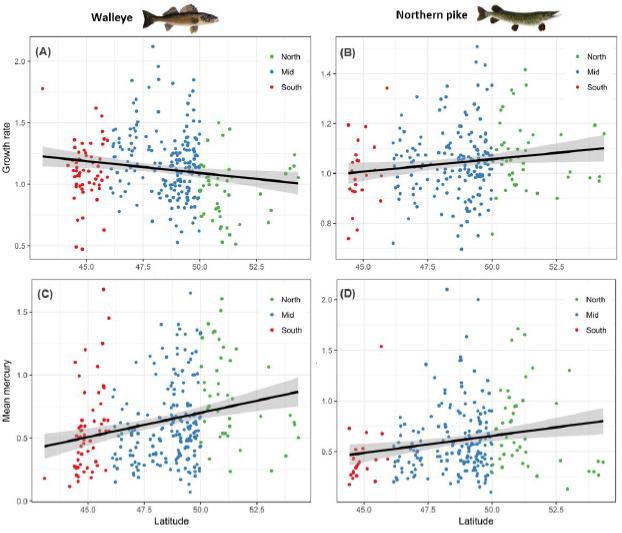
Latitudinal variation in growth rate of A) Walleye (N_Lat_ = 291; R = -0.145; p-value = 0.012) and B) Northern Pike (N_Lat_ = 240; R = 0.125; p-value = 0.05) expressed as random slopes of LMEMs of fish growth with each unique latitude as the random effect, and latitudinal variation in mercury for C) Walleye (N_Lat_ = 291; R = 0.174; p-value = 0.003) and D) Northern pike (N_Lat_ = 240; R = 0.193; p-value = 0.002) expressed as the average amount of fish mercury concentration sampled at a given latitude

### Test of growth dilution

Correlation analysis of mean mercury against random slopes describing growth rates showed an overall negative relationship for Walleye suggesting growth dilution (Supplementary Figure S2-A). In sharp contrast, Northern Pike showed no evidence of growth dilution, and instead revealed a positive relationship between mean mercury and growth rates (Supplementary Figure S2-B). However, the correlations were not significant and correlation coefficients were very low for both Walleye (Pearson’s r = -0.06, p-value =0.33) and Northern Pike (Pearson’s r = 0.11, p-value = 0.08), which implies variation in growth rates fails to explain variation in mercury levels in either species. In summary, the space-for-time substitution approach revealed contrasting growth rate-temperature (i.e. latitudes) relationships between Walleye and Northern Pike, however there is substantial variation in the estimates of both growth rates and mean mercury levels at a given latitude (i.e. temperature), which resulted in the poor correlations between growth rates and mercury levels.

## Discussion

In our analyses of fish mercury levels using LMEMs, the best fitting model was comprised of spatial, temporal and spatiotemporal interactions as random effects; and among these factors, spatiotemporal interactions captured much of the variation. It is thus evident that mercury bioaccumulation in both Walleye and Northern Pike not only varies spatially across Ontario’s freshwater lakes and waterbodies, but is also contingent on the time period. The bioaccumulation index for both Walleye and Northern Pike exhibit fairly complex spatiotemporal patterns across the 40-year time period and three latitudinal zones. There were interesting similarities such as both species showed overall increasing bioaccumulation trends for north and mid latitudes, whereas south latitudes revealed a decreasing trend. There was also substantial variation among the 5–year time periods, as evidenced by the low R-square values for most latitudinal zones, except for Northern Pike in the north and Walleye in the south (Figure 3a,b). Such temporal variations in fish mercury levels have previously been reported from long-term studies (Chalmers et al. 2011; Monson et al. 2011; Gandhi et al. 2014). However, unlike these previous studies, which generally show a declining trend in both north and south latitudes, our findings showed increasing trends in north latitudes, as was the case for Northern Pike, and in mid latitudes, as was the case with Walleye. The reason for this difference in fish mercury trends is perhaps because our definition of bioaccumulation index differs substantially from previous studies that typically measure the rate of change in mercury levels over time. Our measure provides an estimate of the magnitude of bioaccumulation given both latitudinal variability in lake location and temporal variability. Moreover, in our analyses bioaccumulation index is based on the full range of fish size to mercury concentration covariation. Interestingly, like several other long-term studies (Monson et al. 2011; Sadraddini et al. 2011; Gandhi et al. 2014; Blucacks-Richards et al. 2017), our results also show an increase in our mercury bioaccumulation index in north and mid latitudinal zones for the most recent time periods (Figure 3a,b). And this recent increase in fish mercury levels is speculated to be climate change induced (Gandhi et al. 2014). However, as we shall soon discuss, a comprehensive picture of bioaccumulation trends in the context of climate change, suggests complex dynamics that are not easy to generalize.

From a climate change perspective, temperature and growing degree-days showed consistently increasing trends across all latitudinal zones over the 45-year period, while in comparison, mercury bioaccumulation trends surprisingly varied substantially among the latitudinal zones for both Walleye and Northern pike. We hypothesized that the magnitude of growth dilution would be positively correlated with rate of warming, such that latitudinal zones with the highest rates of warming (i.e., the north) would relate to the greatest decrease in fish mercury bioaccumulation. Only Walleye, showed clear evidence of a decreasing trend in mercury bioaccumulation, and this was restricted to the southern latitudinal zone, where relatively slower rates of warming were recorded. It may also be noted that estimated rates of warming did not vary significantly among the three latitudinal zones-though this is perhaps a result of the limited availability of long-term weather data in mid and north latitudinal zones. The implication of nearly similar rates of rising temperature, however, is that temperature driven growth dilution (if present) should be consistent across latitudinal zones, and therefore it is surprising that the latitudinal trends in mercury bioaccumulation do not show the same general pattern. This lack of a general pattern suggests that confounding ecological factors, such as methylation, may play a role. Specifically stated, methylation rates may have increased lately, especially in northern latitudes, since methylation in cold northern latitudes is known to occur during the ice-free season when the soil and ground are not frozen (Stanley et al. 2002; Stern et al. 2012), and northern latitudes are experiencing a more prolonged ice-free season compared to southern latitudes as a consequence of warming climate (Schindler et al. 1990; Dugay et al. 2006). Thus the observed increasing trends in fish mercury levels in the relatively colder north and mid latitudes is possibly due to increasing methylation rates in higher latitudes.

Besides temperature, precipitation is known to affect amount of methylmercury in lakes and waterbodies via surface run-off from surrounding catchment areas (Rudd 1995). Precipitation in Ontario showed a clear increasing trend in time for all three latitudinal zones with northern latitudes showing the greatest rate of increase in time. Such increased precipitation, especially brief intense periods of rainfall can result in enhanced methylmercury levels in lakes (Matilainen et al. 2001; Balogh et al. 2006). Thus, when one considers combined effects of both longer ice-free season and increased precipitation, it is quite apparent that lakes and waterbodies in northern latitudes are likely to end up with greater amounts of methylmercury and consequently higher fish mercury levels over time. In short, the observed latitudinal bioaccumulation trends are due to a complex set of factors that go beyond growth dilution alone.

Unlike spatiotemporal trends of mercury bioaccumulation, the temporal trends estimated using time period as the only random effect showed contrasting species-specific patterns. Northern Pike had consistently increasing trends while Walleye showed a generally declining trend (Figure 3c). It was hypothesized that the contrasting patterns in mercury bioaccumulation over time is driven by differential response of growth dilution to warming temperature, which in turn is driven by temperature-induced variation in growth rate. But the space-for-time substitution analysis (Figure 2) based on growth rate and mean mercury estimates across a latitudinal gradient failed to show significant evidence of temperature-induced difference in growth dilution (Supplementary figure 2). In short, the contrasting bioaccumulation temporal trends in Walleye and Northern Pike are not definitively due to differential response of growth dilution to warming temperature. Evidence for growth dilution from observational studies vary a lot (Karimi et al. 2010), and among the few studies that have reported growth dilution in both Walleye and Northern Pike (Simoneau et al. 2005; Lavigne et al. 2010), growth dilution was inferred from age and mercury estimates of standardized fish lengths. Thus, our results are perhaps partly due to the inclusion of the entire range of body size variation, which adds more variability to the growth rate-mercury covariation compared to standardized fish lengths. Nonetheless, the latitudinal variation in growth rates and mercury levels showed interesting results, which deserve further discussion.

Walleye showed a negative latitudinal growth rate and positive latitudinal correlation with bioaccumulation suggesting growth rates are likely to increase while bioaccumulation decreases with increasing temperature. Previous studies have similarly shown a negative relationship between latitude and growth rates in walleye (Quist et al. 2003; Lavigne et al. 2010). In sharp contrast, for Northern Pike, both growth rate and mean mercury levels showed weak positive correlation with latitude, which suggests that for Northern Pike both growth rate and mercury bioaccumulation are likely to increase with decreasing ambient temperature. Our findings differ from a previous broad-scale analysis of Northern Pike data, where rate of growth was negatively correlated with latitude (Rypel 2012). Moreover, Rypel’s study also reports the presence of a strong counter gradient in growth variation (i.e. potential to adapt growth rate in response to variation in growing period length) when length-at-age was normalized by growing degree-days, suggesting growth in circumpolar fish like Northern Pike is highly variable, and growth rates can potentially increase at higher latitudes to make up for the reduced growing season.

Latitudinal trends in growth rate and mercury levels, however, do not convey the full picture as both Walleye and Northern Pike showed substantial variation in growth rate and mercury levels for any given latitude. This is evident from the low correlation coefficients associated with latitudinal variation in growth rate and mercury levels, suggesting local factors operating at the lake-level have a stronger influence compared to latitude-specific temperature conditions. In a previous study by Simoneau et al. (2005), Walleye populations from different lakes showed evidence of growth dilution, however this was largely driven by lake-specific variation in growth rates. Similarly, a study on mercury contamination in Northern Pike from 19 Boreal lakes showed a high degree of inter-lake variation that was due to lake-specific variations in water chemistry, prey mercury contamination, and landscape-level disturbances in surrounding catchment areas (Garcia and Carignan 2000). In light of these findings, it is perhaps not surprising that strong latitudinal variation in growth rates and mean mercury levels translates into very weak temperature-driven growth dilution effects. This is evident in Walleye, where the space-for-time substitution results suggested that growth rates can potentially increase with temperature, and mercury levels tend to decrease with temperature. These opposing patterns do not translate into a consistent growth dilution effect with warming temperatures due to the presence of a high degree of latitudinal variation in both growth rates and mercury levels in Walleye, thus resulting in the observed weak growth dilution.

Unlike Walleye, Northern Pike showed the opposite of growth dilution with an overall slightly positive correlation between growth rates and mercury levels (Supplementary Figure S2). The weak presence of growth dilution in Walleye (suggestive negative trend) compared to a marginally significant positive trend in Northern Pike is worth noting as it suggests growth magnification may play a modulatory role in Northern Pike’s mercury levels. It is not entirely clear how increase in growth rate results in higher mercury levels in Northern Pike, but it is possible that consumption of highly contaminated food can result in the disproportional addition of mercury relative to biomass. Differences in feeding habits, habitats, and predatory behavior might explain some of the observed difference between Walleye and Northern Pike growth rate-mercury correlation. These differences in feeding ecology might also explain the observed contrasting mercury bioaccumulation trends in time between Walleye and Northern Pike. Studies have reported such disparity between Walleye and Northern Pike in mercury loads as a result of dissimilarities in the feeding habits of these two co-occurring fishes (Mathers et al 1985; Wren et al.1991).

Our study demonstrates how a simple bioaccumulation index derived from mixed-effects models can capture broad-scale patterns of fish mercury bioaccumulation. The index reveals the complex nature of mercury bioaccumulation when both spatial and temporal variations are combined (i.e. spatiotemporal trend) relative to the purely temporal trend. Furthermore, the science and application of ecological indicators now increasingly point to limitations in capturing the complexity of environmental systems and ecosystem responses to various anthropogenic stressors using a single indicator species (Carignan and Villard 2002; Siddig et al. 2016). Hence, it is particularly interesting that the temporal trends captured by the bioaccumulation index highlights strong differences between two co-occurring indicator species in mercury bioaccumulation. And finally, studies on fish mercury bioaccumulation have often stressed growth dilution as a key modulatory mechanism, however the potential of growth dilution to affect large-scale fish mercury dynamics has not been explicitly tested so far. In this respect, our study also shows for the first time that temperature-driven growth dilution has very weak modulatory effect on broad-scale mercury bioaccumulation patterns. From a climate change perspective, this implies change in fish mercury levels as consequence of warming climate is a complex process that goes beyond temperature-driven growth dilution effect.

## Acknowledgements

We are grateful to Tom Johnston, Rob Mackereth, Cindy Chu, Bailey McMeans, and Claire Oswald for feedback during the early stages of the project.

---

**Figure S1.**
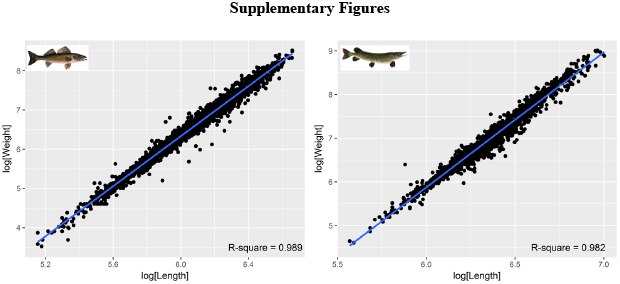
Correlation between fish body mass and body length in Walleye (left) and Northern Pike (right).

**Figure S2.**
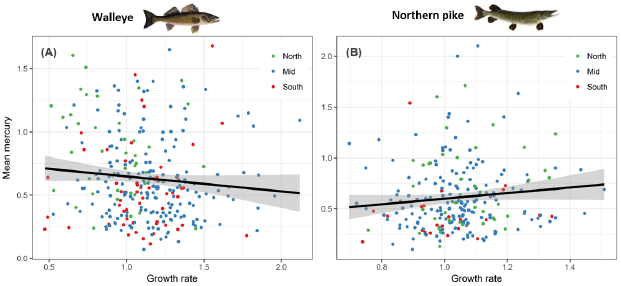
Relationship between estimated growth rates and mean mercury in A) Walleye (NLat = 291; R = -0.06; p-value = 0.33) and B) Northern Pike (NLat = 240; R = 0.11; p-value = 0.08) where mean mercury concentration are based on all samples at a given latitude and growth rates are random slopes of LMEMs of fish growth model with latitude as random effect.

